# Environmental hypoxia controls the evolution of cavefish heart asymmetry

**DOI:** 10.64898/2026.01.29.702597

**Authors:** Mandy Ng, Helena Bilandžija, William R. Jeffery

## Abstract

The environmental factors responsible for the evolution of novel traits and their underlying mechanisms are poorly understood. Here we use the teleost *Astyanax mexicanus* consisting of ancestral surface fish and derived cavefish morphotypes to address the evolution of heart asymmetry. As in other vertebrates, surface fish develop right(D)-looping heart tubes, whereas high levels of left(L)-looping heart tubes have uniquely evolved in cavefish. Cavefish cardiac L-looping is mediated by the upregulation of Sonic Hedgehog (Shh) signalling during gastrulation, which disrupts left-right organizer (LRO) function and modifies the downstream left-oriented Nodal signalling cascade. As a proxy for the original cave colonizers, we exposed contemporary surface fish to key cave-associated environmental factors, namely complete darkness, reduced electrical conductivity, low temperature, hypoxia, or combinations of these conditions. Only hypoxia induced the cavefish L-looping phenotype. Hypoxia increased expression of genes in an integrated Hypoxia Inducing Factor 1(HIF1)-Shh axis, disrupted LRO function, and modified the L-R determination program in surface fish similarly to naturally evolved changes in cavefish. In contrast to surface fish, which show high plasticity in response to hypoxia, cavefish were unable to survive acute hypoxia, suggesting evolutionary refinement and canalization of the ancestral hypoxia response. These studies establish hypoxia as the driver of heart asymmetry and reveal a mechanistic connection between the hypoxic cave environment and the evolution of a novel trait.

## Main

Novel traits evolve as effects of ecological changes on development^1^ but the driving factors and mechanistic links between the environment and new traits are poorly understood. These gaps can be addressed by studying the relatively simple subterranean ecosystem^2, 3^ in which environmental factors and the novel traits in obligate cave-dwelling animals are well known^4^. *Astyanax mexicanus*, a teleost with surface-dwelling (surface fish) and cave-dwelling (cavefish) morphotypes, is an excellent model system for studying phenotypic evolution^5, 6^. Cavefish show reductions or loss of eyes and pigmentation, gains in non-visual sensory organs, the hematopoietic system, and fat reserves, and changes in physiology and behavior^6-8^, which have evolved in a remarkably short evolutionary time frame^9, 10^. One of the most unusual cavefish traits is an alteration in left-right (L-R) heart asymmetry^11, 12^. In surface fish the developing cardiac tube loops to the right of the midline (D-looping) and hearts are formed with a bias toward the left side of the body, a trait conserved throughout the vertebrates ^13-15^. In contrast, cavefish have evolved high levels of cardiac looping to the left of the midline (L-looping) and heart development biased toward the right side of the body^11, 12^.

Visceral organ asymmetry is controlled by the L-R organizer (LRO), known as Kupffer’s vesicle (KV) in teleosts^16^, a small ciliated organ formed at the posterior embryonic midline by the coalescence and differentiation of the dorsal forerunner cells (DFC)^17, 18^. L-R determination is controlled by extracellular fluid flow generated by regional cell shape changes and leftward ciliary beat in the KV^14, 19^. Downstream of ciliary activity, expression of the Nodal antagonist Dand5 on the right but not the left side of the KV activates the Nodal-Pitx2 signalling cascade exclusively in the left lateral mesoderm, which establishes the pattern of visceral organ asymmetry^14^. In cavefish, upregulation of the Sonic Hedgehog (Shh) signalling system along the posterior midline modifies DFC and KV development, resulting in disruption of the downstream Dand5-Nodal cascade and randomization of heart asymmetry^12^. Shh overexpression also contributes to eye degeneration, changes in brain development, and enhancements in the olfactory, gustatory, and erythropoietic systems in cavefish^20-23^.

Despite extensive studies of cavefish development and evolution^6-8^ and knowledge of key environmental pressures that could influence trait evolution in caves^24-27^, namely the absence of light, lack of photosynthesis leading to oxygen reduction (hypoxia), stable temperatures, and decreased electrical conductivity, the drivers of cavefish evolution have not been identified. A notable exception is darkness, which can induce plastic changes in some morphological, metabolic, and neural traits in surface fish that resemble cavefish traits^28^. Phenotypic plasticity may be important for colonization of new habitats^29, 30^, including caves ^28, 31, 32^, and coupled with genetic assimilation and natural selection could facilitate rapid evolutionary changes.

### Hypoxia induces L-looping heart asymmetry in surface fish

Using contemporary surface fish as a proxy for the original cave colonizers, we surveyed the effects of complete darkness, low electrical conductivity, different temperatures, and hypoxia^24-27^ on heart asymmetry. Surface fish embryos, which normally develop >95% D-looping cardiac tubes^11, 12^ (Fig. 1A), were exposed to one or more of these factors beginning at the late blastula stage, and effects on the direction of cardiac looping were assayed at 60 hours post-fertilization (hpf), when the cardiac tube begins to show asymmetry (Fig. 1). To determine if darkness, the unifying factor of all subterranean habitats^3^ and a known inducer of some cavefish traits in surface fish ^28^, affects the development of heart asymmetry, adult surface fish were maintained in the absence of light for about 3 years, and when these fish spawned, their progeny were raised in the dark, and cardiac looping was compared to control embryos spawned from parents maintained under the standard photoperiod. The proportion of D-looping cardiac tubes did not differ between embryos raised in darkness and controls raised in light (Fig. 1E), indicating that darkness does not induce L-looping heart asymmetry. Reduced electrical conductivities are encountered in cave habitats^24-27^, opening the possibility that bioelectrical gradients^33^ may drive heart asymmetry. The effects of conductivity on heart asymmetry were explored in surface fish embryos raised at different conductivity levels. No differences in D-cardiac looping were seen between embryos raised at 690 μS, the conductivity used in *Astyanax* laboratory husbandry^34^, and within the range of water conductivities recorded in surface habitats near caves, compared to 230 μS, within the conductivity range in caves^24, 27^, and at 110 μS, the lowest conductivity tested (Fig. 1F), indicating that low conductivity does not induce cardiac L-looping. Annual temperatures are lower and more uniform in caves than in surface habitats^25-27^, and L-R laterality is influenced by temperature in zebrafish^35^. The possibility that thermal differences may drive heart asymmetry was determined by assaying cardiac looping in surface fish embryos raised at different temperatures. The results indicated that embryos raised at 21°C, 23°C, 26°C, or 30°C showed no differences in the proportion of cardiac L-looping, although the levels of non-looping cardiac tubes were higher at 26°C and 30°C than at lower temperatures (Fig. 1G). In the absence of photosynthesis, cave waters show reduced levels of oxygen compared to surface environments^24, 25, 27, 36^. The effects of hypoxia on heart asymmetry, were determined by exposing embryos to 1 mg/L oxygen in a hypoxia chamber for 4, 8, 15, or 24 hours (hrs) (Fig. 1A-D). We found that L-cardiac looping was increased as a function of hypoxia duration compared to normoxic controls, reaching about 20% after 8 hrs of hypoxia, and 30-40% after 15 or 24 hrs of hypoxia (Fig. 1B-D, H). Notably, L-looping did not exceed 50%, consistent with the randomization of heart asymmetry in cavefish^12^. An increase in the level of non-looping cardiac tubes, another characteristic of the natural cavefish heart asymmetry phenotype^11^, was also evident after 24 hrs of hypoxia (Fig. 1H).

**Figure 1.**
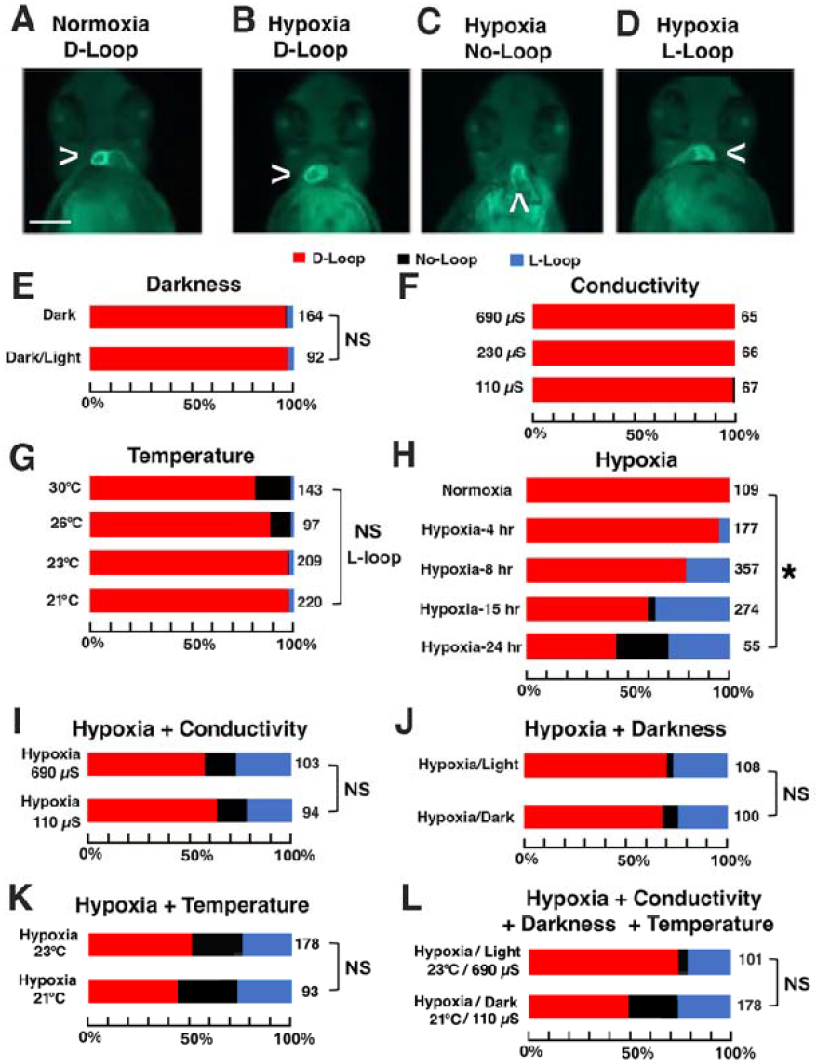
The effects of cave-related environmental factors on heart asymmetry in surface fish. A-D. The direction of cardiac looping in normoxic (A) or hypoxic (B-D) surface fish embryos stained with M-20 myosin heavy-chain antibody. <: D-looping cardiac tube. ^: non-looping (straight) cardiac tube. >: L-looping cardiac tube. Scale bar in A: 250 μm. E-L. The percentage of D-looping (A, B), non-looping (C), and L-looping (D) cardiac tubes in embryos raised in (E) complete darkness versus the standard photoperiod, (F) 690 μS, 230 μS, or 110 μS electrical conductivity, (G) 21°C, 23°C, 26°C or 30°C, (H) normoxia versus hypoxia for 4, 8, 15, or 24 hrs, (I) hypoxia and 690 μS or 110 μS conductivity, (J) hypoxia and standard conditions or complete darkness, (K) hypoxia at 23°C or 21°C, and (L) hypoxia, high conductivity, standard photoperiod at 25°C (control) or hypoxia in darkness, low conductivity (110 μS), and 21°C. Hypoxia exposure began at late blastula, and cardiac looping was assayed at 60 hours post-fertilization. The number of larvae assayed is on the right in each bar. E-L. Statistics by Chi^2^ test. Asterisk in H: Chi^2^ statistic = 120.9265; *p* < .00001. NS: no significance.

Given that ancestral surface fish may have encountered some or all of the environmental pressures described above simultaneously during cave colonization, further experiments explored combinations of hypoxia with one or more of the other cave-related factors. Surface fish embryos were exposed to hypoxia along with (1) the lowest electrical conductivity (110 μS) (Fig. 1I), (2) complete darkness (Fig. 1J), (3) low temperature (21°C) (Fig. 1K), or (4) low conductivity, complete darkness, and low temperature (Fig. 1L), and cardiac looping was compared with controls (Fig. 1I-L). The levels of cardiac L-cardiac looping under conditions 1-4 (above) did not differ significantly from those observed with hypoxia alone, although the proportion of non-looping cardiac tubes was increased in some of these combinations (Fig. 1H-L). Therefore, hypoxia is the only cave-related environmental factor that can change heart asymmetry in surface fish to match the naturally evolved phenotype in cavefish^11, 12^.

### Hypoxia affects left-right organizer development and function in surface fish

We next asked whether hypoxia affects heart asymmetry in surface fish via the same mechanisms that underlie this trait in cavefish^12^. Cardiac L-looping in cavefish is linked to Shh induced increases in DFC leading to changes in KV structure and function and expression of the downstream Nodal signal^12^. Accordingly, we exposed surface fish embryos to hypoxia for 4-6 hr intervals between 3 and 17 hpf, then returned them to normoxic conditions and assayed for cardiac looping at 60 hpf. The results revealed a maximum hypoxia sensitivity period (HSP) between 75% and 90% epiboly (Fig. 2Aa-e), which corresponded to DFC migration and merger into the KV^12^. The effects of hypoxia on DFC number in surface fish were determined by staining with the DFC-specific marker SYTO-11 ^12, 17^, and hypoxic embryos showed more DFC than normoxic controls (Fig. 2B-E, J). Examination of KV morphology showed that 40-50% of the hypoxic embryos formed a major and several minor KVs (Fig. 2K, L), and when single KVs were present they were on average larger than those in normoxic controls (Fig. 2J). The effects of hypoxia on KV function were determined by examining the expression of *dand5* in the KV and the teleost Nodal gene *southpaw* (*spaw*)^37^ in the lateral mesoderm. In contrast to normoxic surface fish embryos, which showed stronger *dand5* expression on the right than on the left side of the KV and *spaw* expression exclusively in the left lateral mesoderm^11-12^, about 40-50% of the hypoxic embryos showed approximately the same *dand5* expression on both sides of the KV or stronger expression on the left than on the right side of the KV (Fig. 2M, N). The extent of modified *dand5* expression in hypoxic embryos (Fig. 2M) was consistent with the proportion of embryos that developed abnormal KVs (Fig. 2L). Furthermore, although *spaw* was expressed in the left lateral mesoderm of normoxic controls (Fig. 2P, Q), *spaw* expression was seen in either the left or right lateral mesoderm, bilaterally, or was entirely absent in hypoxic embryos (Fig. 2P, R). Together, these results show that hypoxia evoked changes in the L-R determination program in surface fish similar to those underlying the L-looping heart phenotype in normoxic cavefish^12^.

**Figure 2.**
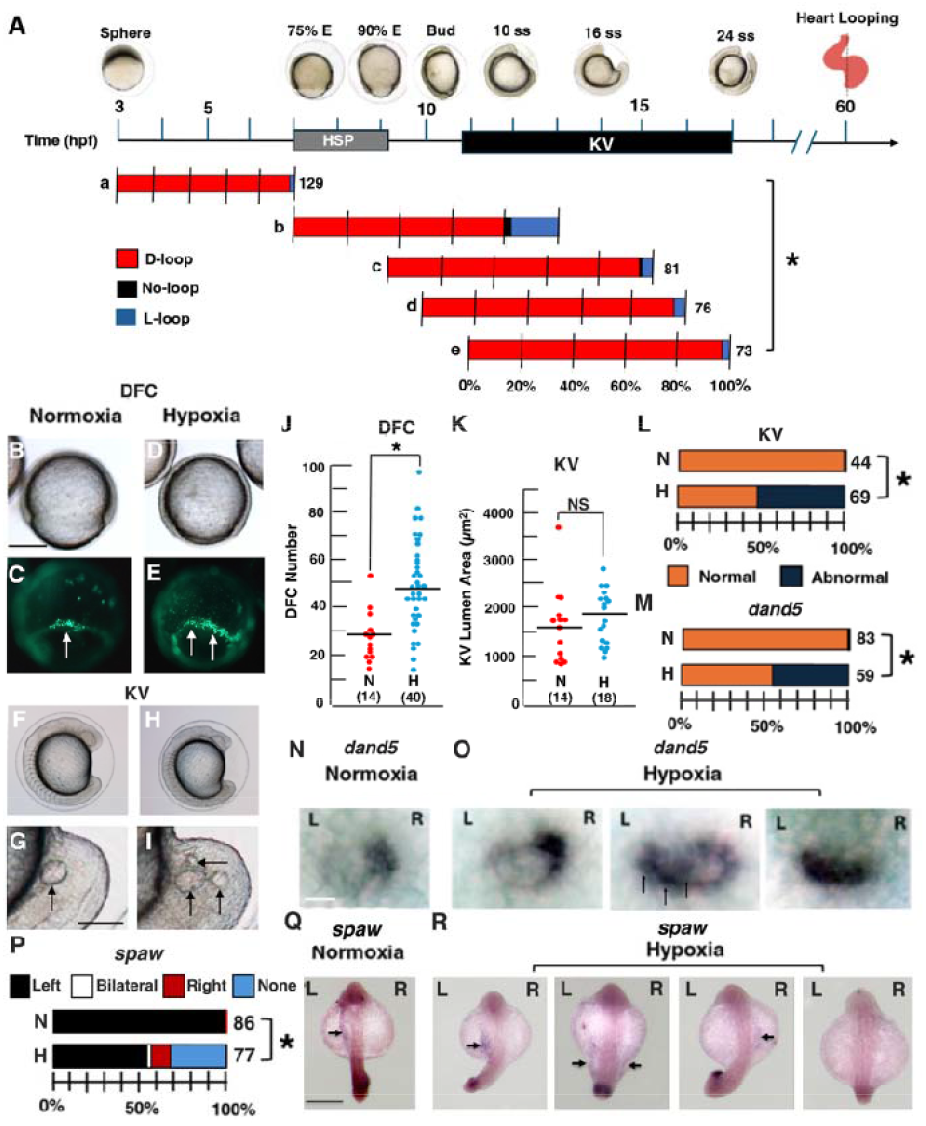
Hypoxia affects left-right organizer development and function in surface fish. A. The hypoxia sensitive period (HSP) for heart asymmetry in surface fish. KV: Kupffer’s vesicle. E: epiboly. Bud: Tailbud. ss: Somites. a-e. Effects of 4 (a) or 6 (b-e) hours of hypoxia on cardiac looping. Statistics: Chi^2^ test. Asterisk: Significance. Chi^2^ statistic = 27.6936. *p* =.000536. B-O. Effects of hypoxia on dorsal forerunner cell (DFC) number (B-E, J), KV development (F-I, K, L), *dand5* expression in the KV (M-O), and *spaw* expression in the lateral mesoderm (P-R). B-I. Comparison of DFC at 70% epiboly by Syto-11 staining (B-E) and KV size at 13-15 ss (E-I) in normoxic (B, C, F, G) and hypoxic (D, E, H, I) embryos. B, D, E-I bright field. C, E. Fluorescence. Arrows in C, E: DFC. Arrows in G, I: KV(s). Scale bar in B: 100 μm. Scale bar in G: 30 μm. Quantification of DFC at 80-90% epiboly (J) and KV lumen areas at 13-15 ss (K) in normoxic (N) and hypoxic (H) embryos. Horizontal lines: means. Number of embryos in parentheses. Asterisk in J: *p* =.000114. NS: No significance. Statistics by One Way ANOVA with Tukey HSD. L. KV with normal and or abnormal morphology (see I) in normoxic (N) and hypoxic embryos (H) at 13-15 ss. Number of embryos shown on the right. Statistics by Chi^2^ test. Chi^2^ statistic: 21.9921. Asterisk: *p* <.00001. M-O. Effects of hypoxia on asymmetric *dand5* mRNA expression. N, O. In situ hybridization showing normal *dand5* asymmetry with (N) expression restricted to the right side of the KV in normoxic embryos and (O) abnormal *dand5* asymmetry in hypoxic embryos at the 13-15 ss stage. M. The proportion of normoxic (N) and hypoxic (H) embryos at the 13-15 ss stage with normal or abnormal *dand5* expression asymmetry. Number of embryos shown in the right of bars. Statistics: Chi^2^ test. Chi^2^ statistic: 41.9981. Asterisk: Significance at *p* <.00001. Q-R. In situ hybridization showing *spaw* expression (arrows) in lateral mesoderm of normoxic embryos and hypoxic embryos. P. Summary of normoxic (N) and hypoxic (H) embryos at the 25 ss stage with normal left-oriented or abnormal *spaw* expression in the lateral mesoderm. Number of embryos shown to the right of each bar. Statistics by Chi^2^ test. Chi^2^ statistic: 34.9535. Asterisk: Significance at *p* <.00001. In N, O, Q, R: L is left and R is right. Hypoxia for 8 hrs. beginning at late blastula in B-R.

### Hypoxia upregulates the HIF1α-Shh axis in *Astyanax*

As a response to hypoxia, the transcription factor HIF1α is stabilized and accumulates in nuclei, upregulating multiple downstream targets^38-40^ including the Shh signalling pathway^23, 41-46^. Importantly, the increased Shh pathway is also linked to the evolution of heart asymmetry^12^ and many other traits in cavefish^20-23^. Therefore, we asked whether hypoxia affects expression of the HIF1α-Shh axis (Fig. 3). Embryos were exposed to hypoxia and expression of the HIF1α target genes *erythropoietin a* (*epoa), glucose-6-phosphate dehydrogenase* (*gapdh*), *hexokinase 1* (*hk-1*), and *insulin-like growth factor binding protein 1a* (*igfbp1a*)^38, 47-52^, the paralogous *shha* and *shhb* genes, and the Shh-dependent *gli1* and *foxj1a* genes^12, 53^ was compared to normoxic controls by qRT-PCR. Hypoxia applied during HSP increased expression of most of these HIF1α-Shh axis genes in surface fish and cavefish by about 2-fold or more relative to normoxic controls (Fig. 3A; Table S1). We investigated the relationship between hypoxia and the HIF1α-Shh axis using cell-permeable small molecule antagonists and agonists: echinomycin, which selectively inhibits HIF1α transcriptional activity by preferential binding to the core DNA recognition element^54, 55^, DMOG, which activates HIF1α by inhibiting prolyl-hyroxylase activity under normoxic conditions^56-58^, and the Smoothened inhibitor SANT1^59^, which reduces Shh signaling in *Astyanax* embryos^12^. Co-exposure of surface fish embryos to echinomycin and hypoxia decreased the proportion of cardiac L-looping relative to controls exposed only to hypoxia (Fig. 3B). Conversely, treatment of surface fish with DMOG significantly increased cardiac L-looping in the absence of hypoxia (Fig. 4C). Lastly, SANT1 treatment during hypoxia reduced L-cardiac looping in surface fish compared to controls exposed only to hypoxia (Fig. 3D), and co-exposure to echinomycin and hypoxia together compared to hypoxia alone narrowed the *shha* expression domain along the embryonic midline (Fig. 3E-G), confirming the link between the hypoxia-induced HIF1α and Shh signalling pathways.

**Figure 3.**
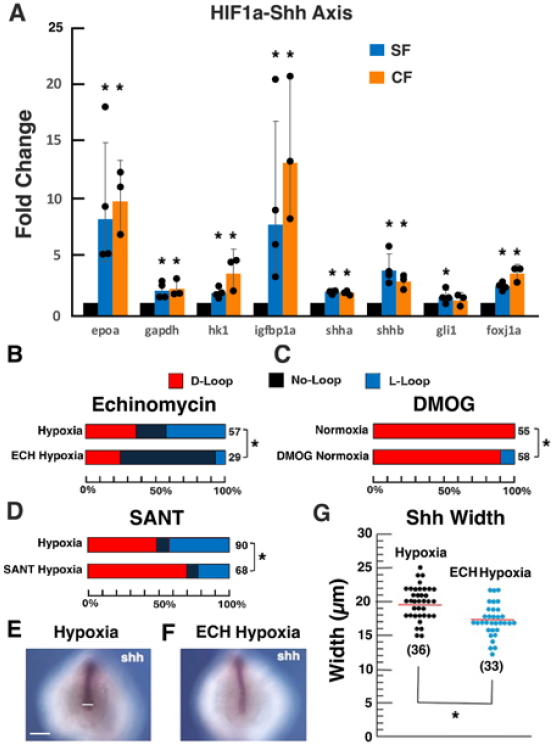
Role of the HIF1α-Shh axis in hypoxia-induced heart asymmetry. A. HIF1α-Shh axis gene expression quantified by qRT-PCR with at least 3 biological replicates in 13-15 somite hypoxic (15 hrs beginning at late blastula) versus normoxic surface fish and cavefish embryos. Asterisks (Significance; See Table S1). SF: surface fish. CF: cavefish. B-D. Surface fish embryos treated with (B) echinomycin and hypoxia (bottom) compared to hypoxia alone (top), (C) DMOG (bottom) compared to normoxia control (top), and (D) SANT-1 and hypoxia (bottom) and hypoxia (top) compared to hypoxia control. Numbers of larvae shown at the right. Statistics by Chi^2^ test. Asterisks: B, Chi^2^ statistic = 18.005, *p* = 000123; C, Chi^2^ statistic = 6.6204, *p* =.036509; D, Chi^2^ statistic = 7.3631, *p* =.025184. E, F. In situ hybridization showing *shha* expression at the posterior midline in 13-14 somite surface fish embryos exposed to hypoxia (E) or hypoxia and echinomycin (F) for 8 hrs beginning at the late blastula stage. Scale bar; 50 μm; White horizontal line in center of the frame indicates where *shha* expression width was measured in G. G. Summary of *shha* expression width differences at the posterior midline of embryos exposed to hypoxia or hypoxia and echinomycin. Red horizontal lines represent means. Number of embryos in parentheses. Asterisk: p =.000859. Statistics by one way ANOVA with Tukey HSD. ECH: echinomycin. B-G: Hypoxia for 8 hrs beginning at late blastula.

**Figure 4.**
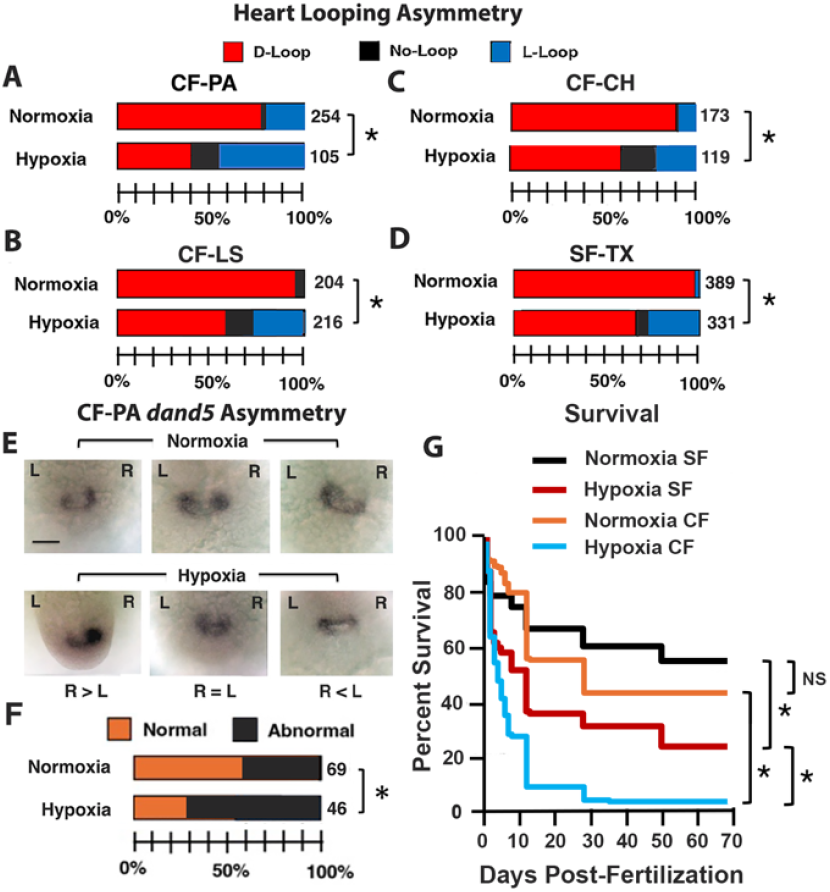
Hypoxia affects heart asymmetry and survival in cavefish. A-D. The effects of hypoxia on heart looping in different cavefish populations and the Texas surface fish compared to normoxic controls. A. Pachon cavefish (CF-PA). B. Los Sabinos cavefish (CF-LS). C. Chica cavefish (CF-CH). D. Texas surface fish (SF-TX). The number of embryos shown on the right. Hypoxia for 15 hrs beginning at late blastula. Asterisks: Significance by Chi^2^ tests: (A) Chi^2^ statistic 84.4616; p <.00001, (B) Chi^2^ statistic 82.7638; p <.00001, (C) Chi^2^ statistic 30.2603; p <.00001, and (D) Chi^2^ statistic 114.7925A; p <.00001. E-G. Effects of hypoxia on *dand5* expression asymmetry in the KV of 13-15 somite CF-PA embryos. E. In situ hybridization showing *dand5* expression in normoxic (top) and hypoxic (bottom) CF-PA embryos. F. Summary of E. L: left. R: right. Numbers of embryos on the right. Asterisk: Significance by Chi^2^ test. Chi^2^ statistic 5.7773, p =.022235. H. Survival of surface fish (SF) and cavefish (CF) exposed to hypoxia for 15 hrs beginning at late blastula (Fig. 2A), compared to normoxic controls. N: 76 for normoxic and 108 for hypoxic surface fish, and 180 for normoxic and 165 for hypoxic cavefish. Asterisks: 1: *p* =.000029; 2: *p* <.000001; 3: *p* <.000001. NR: no significance; *p* =.161188. Statistics by Cox regression^66^.

### Effects of hypoxia on heart asymmetry and survival in cavefish

To determine whether the effects of environmental hypoxia on heart asymmetry are ubiquitous in diverse *Astyanax* populations, we exposed three different cavefish populations (Pachón, CF-PA; Los Sabinos, CF-LS; and Chica, CF-CH) and surface fish from Texas (SF-TX), located about 1,500 km from the Mexican population (SF-MX), to hypoxia during HSP (Fig. 2A). The results showed that normoxic PA-CF and CH-CF have elevated cardiac L-looping relative to normoxic LS-CF and both surface fish lineages (Figs. 2H; 4A-D), in line with their origin from two of the most hypoxic of all *Astyanax* caves^27^. Furthermore, hypoxia increased cardiac L-looping in each cavefish population, including those populations that already showing high levels of L-looping under normoxic conditions (Fig. 4A-D), suggesting that hypoxia effects on heart asymmetry may be ubiquitous in *Astyanax*. However, the levels of L-looping induced by hypoxia did not surpass about 50%, consistent with the randomization of heart asymmetry (Fig. 1H). The effects of hypoxia on KV-associated *dand5* expression were determined in PA-CF embryos, and the results were similar to effects on surface fish (Figs. 2N, O, 4E, F), confirming the conclusion that hypoxia drives heart asymmetry by increasing the HIF1α-Shh axis and disrupting LRO function in both surface fish and cavefish.

To understand whether the plastic hypoxia responses are the same in both the ancestral and derived lineages, we compared the survival of hypoxia-exposed surface fish and cavefish through the juvenile stages (Fig. 4G). About 50-60% survival of normoxic surface fish and cavefish was observed after 70 days, consistent with previous results in our culture conditions^11, 60^. After a 15 hr pulse of hypoxia (including HSP), about 25% of surface fish survived for 70 days, but cavefish were more sensitive to hypoxia and showed less than 5% survival after only 30 days (Fig. 4G). These results indicate that surface fish have greater tolerance to hypoxia than cavefish, despite the origin of cavefish in hypoxic caves and their evolved adaptations to low oxygen^23, 52, 61, 62^. Thus, the original hypoxia response induced during cave colonization appears to have been refined and canalized during subsequent cavefish evolution.

## Discussion

This study dissects the environmental components of caves to find the specific ecological factor and mechanism responsible for the evolution of a novel cavefish trait: heart asymmetry. A brief exposure of surface fish to hypoxia during late gastrulation induced 30-50% cardiac L-looping during heart development in surface fish, a trait that is unique to cavefish. In contrast, no changes in heart asymmetry appeared after exposure to complete darkness, reduced electrical conductivity, low temperature, or when one or more of these cave-related factors were combined with hypoxia, representing a more realistic simulation of the natural cave environment. Hypoxia induced changes in heart asymmetry were detected in two diverse surface fish populations and three different cavefish populations, implying that oxygen reduction may be a ubiquitous generator of heart asymmetry changes in *Astyanax*. The existence of hypoxia-induced changes in heart asymmetry in diverse surface fish lineages suggests that this plastic trait is ancestral and evolved before cave colonization, but became especially relevant in caves where behavioral responses could not be used to avoid hypoxic waters^64, 65^. Our results provide strong evidence that environmental hypoxia is the sole factor in the cave environment that can modify the conventional pattern of heart asymmetry, which is conserved in all vertebrates.

The natural L-looping cardiac phenotype of cavefish is linked to changes in the program of L-R asymmetry determination, including upregulation of the Shh signalling pathway, modifications of DFC and KV development, and changes in the asymmetric Dand5-Nodal cascade responsible for establishing the direction of heart asymmetry^12, 15^. The same effects on the molecular and developmental program underpinning heart asymmetry as occur in normoxic cavefish were detected following exposure of surface fish to hypoxia, showing that hypoxia modifies the L-R determination pathway in surface fish to resemble the naturally evolved program in cavefish. However, cardiac L-looping was not increased above 50% by hypoxia in either surface fish or cavefish, suggesting that reduced oxygen causes randomization of heart asymmetry rather than a specific reversal from cardiac D-to L-looping. Together, these results indicate that exposure of surface fish to hypoxia phenocopies the natural developmental mechanisms underlying evolutionary changes in cavefish heart asymmetry and open the possibility that other Shh-dependent cavefish traits, including eye degeneration and constructive traits^19-23^, may have also evolved through upregulation of the hypoxia-induced HIF1α-Shh axis.

Our findings suggest a possible scenario for the colonization of hypoxic cave ecosystems by surface fish ancestors and the subsequent rapid evolution of cavefish. First, the high plasticity of the hypoxia response in surface fish, including increases in hematopoietic domains, erythrocytes, and the respiratory system ^23, 52, 54, 61-63^, may have pre-adapted surface fish colonizers for survival in caves with reduced oxygen. The ability to mate and spawn in complete darkness^28^ would also have been a key factor for establishing reproducing populations in caves. Second, after cave colonization, the linkage between hypoxia, HIF1α stabilization^38-40^, and hyperexpression of the Shh signalling system^38-40^ could have triggered the appearance of constructive traits, such as increases in olfaction and gustation^20-22^, which support survival in the dark cave environment where visual senses are obsolete and would serve as targets of natural selection. Third, cavefish may have refined and canalized the ancestral hypoxia response over subsequent generations because the original plasticity was no longer necessary in the homogeneous cave environment.

In conclusion, since perpetual darkness and the absence of photosynthesis are unifying themes in caves^3^, hypoxia may be a common but previously unappreciated contributor to the evolution of novel traits during adaptation to subterranean environments.

## Methods

### Biological materials and husbandry

*A*styanax *mexicanus* surface fish and cavefish embryos were obtained by spawning of adult stocks in the Jeffery laboratory. Surface fish stocks were originally collected in Nacimiento del Rio Choy, San Luis Potosi, Mexico (SF-MX) and San Solomon Springs, Balmorhea, Texas, USA (SF-TX). Cavefish stocks were originally collected in Pachón (CF-PA) cave in Tamaulipas, Mexico, and Los Sabinos (CF-LS) and Chica (CF-CH) caves in San Luis Potosi, Mexico. Surface fish and cavefish were raised in a constant flow aquatic system under normoxic conditions at 23°C on a 14 hr light and 10 hr dark photoperiod (standard conditions). Spawning was induced by increased feeding and water temperature, embryos and larvae were raised under standard conditions^34^, and embryos were staged according to Hinaux et al. ^67^. Experimental protocols were executed according to University of Maryland, College Park procedures (IACUC #R-NOV-18-59; Project 1241065-1 and #R-FEB-25-04; Project 2242381-1) in compliance with ARRIVE guidelines.

### Raising surface fish adults and embryos in complete darkness

Surface fish adults were raised in complete darkness beginning during the early cleavage stages for about 3 years as described by Bilandžija et al. ^28^, and controls were raised for the same period under standard conditions. Dark-raised surface fish adults spawned spontaneously in the absence of light, and the embryos were raised in complete darkness throughout embryogenesis in aquatic system water (ASW) until 60 hours post-fertilization (hpf), when they were assayed for cardiac looping asymmetry as described below. Control embryos were spawned from adults raised in ASW under standard conditions and assayed for cardiac looping asymmetry at 60 hpf.

### Raising surface fish embryos at different electrical conductivity

Surface fish embryos were raised in ASW under standard conditions at high (normal), medium, or low electrical conductivity for 8 or 15 hrs beginning at the sphere stage (3-4 hpf), subsequently washed into ASW of normal conductivity, and then returned to standard conditions and assayed for cardiac looping at 60 hpf (see below). High (normal) conductivity was 690 μSiemens (μS), the conductivity used for fish husbandry^34^ and within the range of conductivity measured in surface waters near *Astyanax* caves^24, 25, 27^, medium conductivity was 230 μS, the conductivity reported in some *Astyanax* caves^24, 27^, and low conductivity was 130 μS. Water of different conductivity levels was prepared by dilution of 690 μS ASW with appropriate amounts of Millipore Q distilled water (conductivity 18 μS). Conductivity was monitored using a Eutech Elite PTCS probe (Thermo Scientific; Waltham, MA, USA).

### Raising surface fish embryos at different temperatures

Surface fish embryos were raised in ASW at 21°C, 23°C, 26°C, or 30°C for 15 hrs beginning at the sphere stage, then incubated in ASW under standard conditions, and assayed for cardiac looping at 60 hpf.

### Raising surface fish and cavefish embryos exposed to hypoxia

Dissolved oxygen levels between 0.9 and 6.9 mg/L have been recorded in different caves in the Mexican Sierra del Abra region^24, 25, 68, 69^, and these levels are likely to fluctuate seasonally^27^. Therefore, we selected 1 mg/L oxygen, which is slightly above the lowest recorded value, for this study. The low oxygen conditions were applied in a hypoxia chamber (ProOx Model P110, BioSpherix, Parish, NY, USA) by bubbling nitrogen gas (HP, Airgas, Hyattsville, MD, USA) into the aquatic environment. About 50 embryos at the late blastula stage were exposed to hypoxia for 4-24 hrs in glass dishes containing 150 ml of ASW that had been pre-incubated for 6-24 hrs in the hypoxia chamber. Oxygen levels were monitored and adjusted automatically by an oxygen sensor in the hypoxia chamber. Hypoxia exposure was terminated by rinsing embryos into normoxic ASW, and culture was continued under standard conditions. Normoxic control embryos were raised in ASW under standard conditions outside of the hypoxia chamber. After the completion of hypoxia, some embryos and normoxic controls were immediately prepared for RNA extraction or fixed for in situ hybridization at the 13-15 or 25 somite stages, and others were raised until 60 hpf for cardiac looping assays.

### Raising surface fish embryos exposed to combined environmental factors

Surface fish embryos were exposed to hypoxia along with other cave-related environmental factors (complete darkness, 21°C, 230 μS conductivity), either individually or simultaneously, beginning at the late blastula stage and lasting for 8 hrs. After the completion of these treatments, the embryos were cultured in ASW under standard conditions and assayed for cardiac looping at 60 hpf.

### Cardiac looping asymmetry

Cardiac looping asymmetry was assayed in larvae fixed overnight with 4% paraformaldehyde (PFA) and stained with myosin heavy chain MF-20 antibody^70^ (Developmental Studies Hybridoma Bank, University of Iowa, Iowa City, IA, USA) or in living larvae anesthetized with 2 mg/ml MS222 (Tricaine; Western Chemical Inc, Ferndale, CA, USA). Imagining was from the ventral side with a Zeiss Axioskop compound microscope as described by Ma et al.^34^. Embryos were classified as having right (D) looping, no (straight) looping, or left (L) looping cardiac tubes.

### Survival curves

Survival curves were constructed for surface fish and cavefish embryos exposed to hypoxia for 15 hrs beginning at late blastula and normoxic controls. Following hypoxia exposure, the larvae were raised for 69 days under standard conditions in glass bowls containing 50 ml ASW, fed brine shrimp beginning at 6 dpf, and counted periodically under a stereomicroscope after brief anesthetization as described by Ma et al^34^.

Following quantification, the larvae were rinsed several times in ASW and returned to the bowls. Dead larvae were removed from the bowls daily. Statistical significance was determined according to the Cox proportional hazards model^66^.

### Dorsal forerunner cell quantification

Endocytic DFC were labeled with the fluorescent dye SYTO 11 (Invitrogen, Carlsberg, CA, USA) as described by Cooper and D’Amico^17^ and Ng et al.^12^. Surface fish embryos exposed to hypoxia as described above and normoxic controls were dechorionated by treatment with 0.2 mg/ml protease type XIV (Sigma-Aldrich, St. Louis, MO, USA) for 1 min at room temperature, rinsed in ASW, and then incubated in 15 μM SYTO 11 diluted in DMSO. Control embryos were treated with 2% DMSO in ASW. Embryos were imaged for fluorescent DFC quantification at 70-80% epiboly or for KV visualization at the 13-15 somite stage as described by Ng et al.^12^.

### Kupffer’s vesicle measurements

The KVs of 13-15 somite surface fish embryos exposed to hypoxia as described above and normoxic controls were live imaged and photographed with a Zeiss Axioskop microscope. KV lumen perimeters were outlined and converted into areas using ImageJ software as described by Ng et al.^12^.

### Modulation of the HIF1α and Shh signaling pathways

Surface fish embryos were pre-exposed to the HIF1α DNA-binding inhibitor echinomycin (10 μM; Abcam, Cambridge, UK), the HIF1α pathway stabilizer dimethyloxallyl glycine (DMOG) (50 μM, Tocris Bioscience, Bristol, UK), or the Smoothened inhibitor SANT-1 (2 μM, Tocris) for 8 hrs. The pharmacological agents were diluted with DMSO, stored at -20°C, and appropriate DMSO concentrations were used in controls. The embryos were pre-treated with echinomycin or SANT-1 for 1 hrs or 1.5 hrs, respectively, prior to hypoxia exposure. At the end of hypoxia exposure, aliquots of some embryos were removed for RNA isolation and other embryos were rinsed into ASW and raised under standard conditions until 60 hpf for cardiac looping assays.

### In situ hybridization

Surface fish or cavefish embryos were dechorionated by protease treatment (see above) or by manual removal with forceps, then fixed in 4% PFA in phosphate buffered saline (PBS) overnight, dehydrated in increasing methanol concentrations to 100%, and stored at -20 °C. RNA probes for *dand5, shha*, and *spaw* mRNA detection were prepared by RT-PCR from surface fish cDNA using oligonucleotide primers (Table S2) designed from sequence information in the *A. mexicanus* surface fish genome assembly^71^ as described previously^12^. In situ hybridization was carried out as described by Ma et al.^72^. After the completion of hybridization, the embryos were washed with PBS-0.1% Tween and incubated in BM Purple AP Substrate (Roche, Basel, Switzerland) at room temperature in the dark. After the signal developed, the reaction was stopped by rinsing the embryos in PBS. The stained embryos were processed through an increasing glycerol series in PBS and photographed using a Zeiss Axioskop compound microscope. Measurements of the lateral extent of *shha* expression at the posterior midline were done with 13-15 somite embryos as described by Ng et al.^12^.

### RNA isolation and quantitative reverse transcriptase polymerase chain reaction

Surface fish and cavefish embryos were immersed in TRI Reagent Solution (Life Technologies, Grand Island, NY, USA) and total RNA was isolated using the Direct-zol RNA Microprep kit (Zymo Research, Irvine, CA, USA). Total RNA was used in quantitative reverse transcriptase polymerase chain reaction (qRT-PCR) experiments after removal of genomic DNA by treatment with RNase-free DNase I (Zymo Research), and cDNA was synthesized using the SuperScript IV VILO Master Mix. qPCR was performed using the TB Green Premix Ex Taq II (Takara Bio, San Jose, CA, USA) under the following cycling conditions: initial denaturation for 30 sec. at 95°C, then 40 cycles for 5 sec. at 94°C and 30 sec. at 60°C, and finally 5 sec. at 95°C, 1 min. at 60°C, and 95°C in increments of 0.11°C/sec. Sequence information from the *A. mexicanus* surface fish genome assembly^71^ was used to design oligonucleotide primers for qPCR (Table S3). The *rpl11* and *smarce1* genes were used as references. The qRT-PCR reactions were performed using a LightCycler 480 (Roche, Indianapolis, IN, USA). The ΔCt for each gene was calculated by subtracting the average Ct value of the reference genes, and ΔΔCt was calculated by subtracting the average ΔCt of the experimental genes from the control gene group. The fold change was calculated as 2^−(ΔΔCt)^. The ΔΔCt values were used for statistical analyses.

## Acknowledgements

We thank Carinna Householder for conducting pilot studies on the role of hypoxia in heart asymmetry, and Lauren Cox, Karina LaCroix, Ruby Dessiatoun, and Barend Keller for husbandry of the *Astyanax* colony.

## Funding

This research was supported by research funds from the University of Maryland, College Park (WRJ) and a postdoctoral fellowship from FP7 People: Marie-Curie Actions (FP7-PEOPLE-2011-COFUND – NEWFELPRO FP7 2007-2013, grant agreement no. 291823, project EACAA, agreement No. 50) (HB).

## Author Contributions

M. N. conducted the experiments, W. R. J. designed the project, and H. B. and W. R. J. supervised the project and wrote the manuscript.

## Competing Interests

The authors declare no competing interests

## Supplementary Information

**Table S1.** Gene expression fold changes and statistical significances of HIF1α-Shh axis genes.

| Gene | SF H versus N |  | CF H versus N |  |
| --- | --- | --- | --- | --- |
|  | Fold change | p value | Fold change | p value |
| *epoa* | 8.083 | .000112 | 9.586 | .000106 |
| *gapdh* | 2.036 | .002571 | 2.187 | .005325 |
| *hk1* | 1.840 | .007165 | 3.476 | .00492 |
| *igfbp1* | 7.637 | .000043 | 12.835 | .000312 |
| *shha* | 1.948 | .000154 | 1.877 | .000171 |
| *shhb* | 3.759 | <.000001 | 2.808 | .000731 |
| *gli1* | 1.711 | .01282 | 1.222 | .236027 |
| *foxj1a* | 2.428 | <.000001 | 3.477 | .000201 |

Analysis by qRT-PCR in 13-15 somite normoxic (N) and hypoxic (H) surface fish (SF) and cavefish (CF) embryos. Statistics by Students t test.

**Supplementary Table S2.**
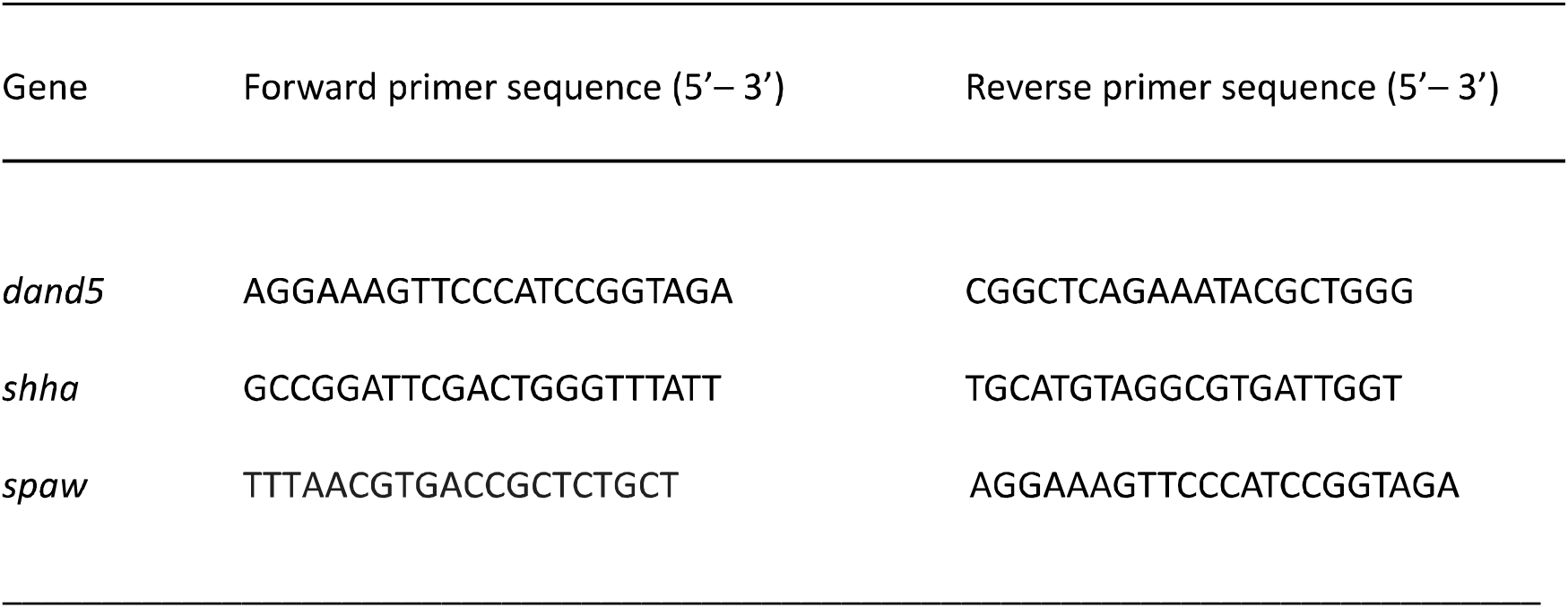
Oligonucleotide primer sequences used generate RNA probes for in situ hybridization.

**Supplementary Table S3.**
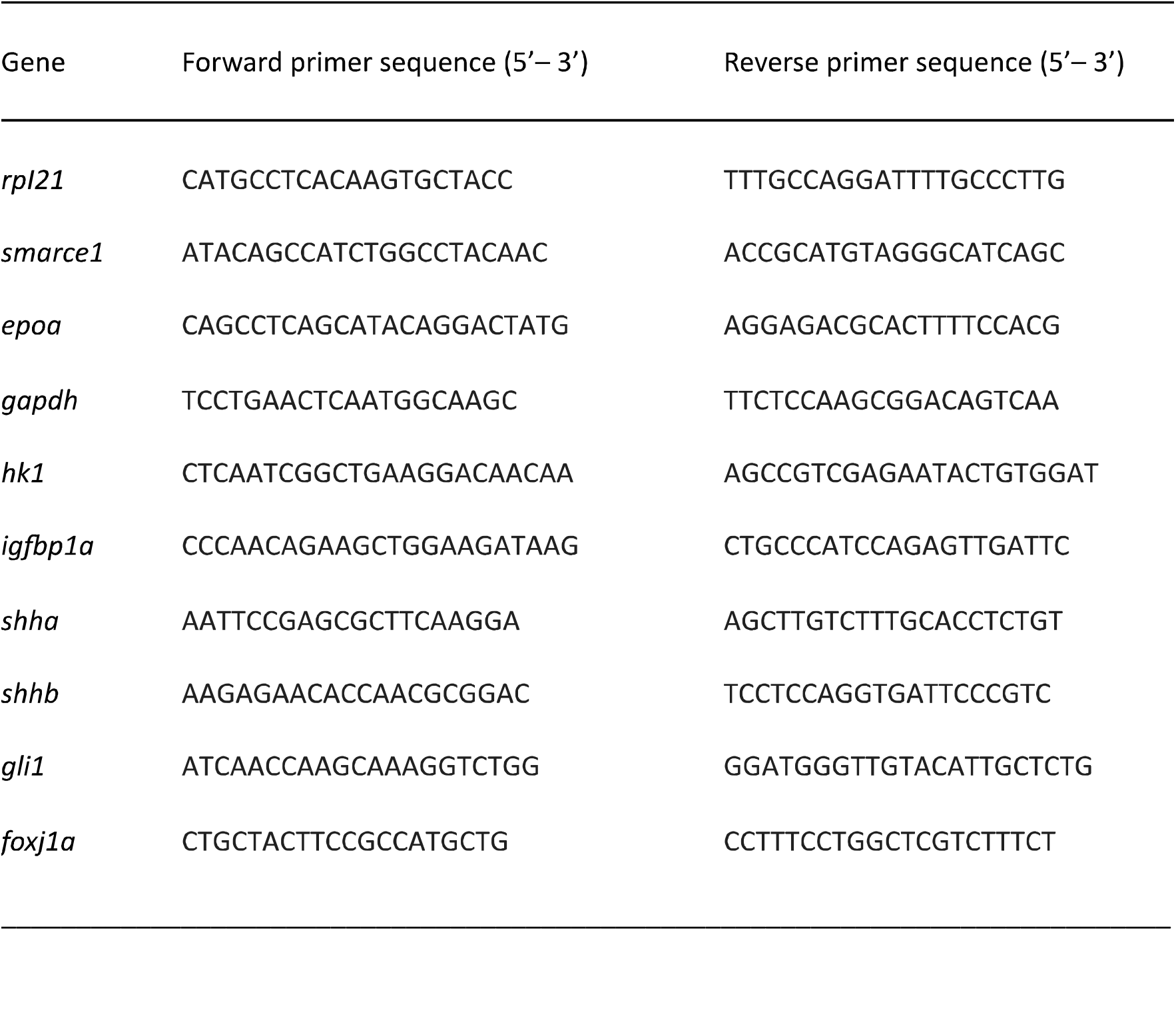
Oligonucleotide primer sequences for qPCR.

## Notes

### Competing Interest Statement

The authors have declared no competing interest.

